# A multifaceted approach to analyzing taxonomic, functional and phylogenetic β-diversity

**DOI:** 10.1101/814467

**Authors:** Gabriel Nakamura, Wagner Vicentin, Yzel Rondon Súarez, Leandro Duarte

## Abstract

Ecological literature offers a myriad of methods for quantifying β-diversity. One such methods is determining BD_total_ (BD), which, unlike other methods, can be decomposed into meaningful components that indicate how unique a community is regarding its composition (local contribution) and how unique a species is regarding its occurrence in the metacommunity (species contribution). Despite this advantage, the original formulation of the BD metric only assesses taxonomic variation and neglects other important dimensions of biodiversity. We expanded the original formulation of BD to capture variation in the functional and phylogenetic dimensions of a metacommunity by computing two new metrics — BD_Fun_ and BD_Phy_ — as well as their respective components that represent the local and species contribution. We tested the statistical performance of these new metrics for capturing variation in functional and phylogenetic composition through simulated metacommunities and illustrated the potential use of these new metrics by analyzing β-diversity of stream fish communities. Our results demonstrated that BD_Phy_ and BD_Fun_ have acceptable type I error and great power to detect the effect of deep evolutionary relationships and attributes mediating patterns of β-diversity. The empirical example illustrates how BD_Phy_ and BD_Fun_ reveal complementary aspects of β-diversity relative to the original BD metric. These new metrics can be used to identify local communities that are of conservation importance because they represent unique functional, phylogenetic and taxonomic compositions. We conclude that BD_Phy_ and BD_Fun_ are important tools for providing complementary information in the investigation of the structure of metacommunities.

## Introduction

Since Whitaker’s seminal paper (Whittaker 1960), many measures and definitions have been proposed to refer to and operationalize β-diversity. Despite the diversity of mathematical formulations and propositions to operationalize this concept (Anderson et al. 2011 for a review), at the core of any β-diversity metric is the notion that it has the purpose of capturing patterns of variation in community composition (Legendre et al. 2005), which in turn can be used to describe ecological patterns that can shed light on the processes and mechanisms that affect the distribution of species in space and time (Baselga 2010).

Among the plethora of methods to measure β-diversity is the framework called BD_total_ (hereafter only BD) proposed by Legendre & De Cáceres (2013), which deserves special attention due to its computational simplicity and the meaningful components that can be extracted from it. Among the advantages of this method we stress its mathematical independence (the ability to compute β-diversity independently of computing alpha and gama diversity) (Ellisson 2010), as well as the possibility to decompose the total variation present in a metacommunity matrix it into two components: Local Contribution to Beta Diversity (LCBD), which indicates the portion of total variation accounted for by an individual sample/community in a metacommunity, and Species Contribution to Beta Diversity (SCBD), which indicates the contribution of individual species to total BD. Whereas BD can be interpreted as a general measure of β-diversity for a metacommunity, LCBD and SCBD represent, respectivelly, how unique communities and species are in that metacommunity (Legendre and De Cáceres, 2013).

Despite the advantages of the BD framework and its components for revealing ecological patterns in community structure (e.g. Li et al. 2019; Yao et al. 2019), it does not consider variation from other dimensions of biodiversity that could shed light on evolutionary and ecological patterns of species distributions across communities. It is known that traditional taxonomic metrics of β-diversity combined with functional and phylogenetic measures can shed light on the balance between environmental and evolutionary factors affecting the composition of communities (Graham and Fine 2008; Pillar et al. 2009; Pillar and Duarte 2010; Duarte et al. 2016; Safi et al., 2011). Among the practical advantages of considering variation in different dimensions of biodiversity is the possibility of identifying sites that concentrate functional, phylogenetic and taxonomic diversity as being of special interest for conservation proposes (Devictor et al. 2010). Consequently, statistical tools capable of characterizing variation in different components of biological diversity are needed for improving our understanding of factors acting on the structure of metacommunities.

Therefore, in order to join the advantages of the BD framework with the possibility of capturing other components of variation in biological diversity, we show herein how BD can be extended to produce β-diversity metrics that represent phylogenetic and functional dimensions of diversity, while preserving all the advantages of the original BD framework. Specifically, our goals were to: (1) expand the BD framework by deriving two new metrics called BD_Phy_ and BD_Fun_ and their respective components of local and species contributions to β-diversity; (2) test the performance of BD_Phy_, BD_Fun_ and their local components in capturing variation in metacommunity composition and community uniqueness mediated by the functional and phylogenetic relationship among species; and (3) show how these metrics can be used together to reveal patterns of variation in metacommunities by using as an example a data base for a tropical stream fish metacommunity.

## Methods

### Expanding the BD framework to include functional and phylogenetic dimensions of biodiversity

There are two ways to compute BD: directly from a matrix of species composition per site, or from a distance matrix calculated from a species composition matrix using an adequate dissimilarity index. For the sake of simplicity, we will demonstrate how BD can be extended using only the procedure involving distance matrices.

Given a matrix **W** describing *i* sites (rows) by *j* species (columns), the first step consists of redescribing the species occurrence matrix **W** to represent the clade and trait distribution of species across communities (Pillar & Duarte, 2010; Duarte et al., 2016). The redescription of matrix **W** to obtain the two new matrices starts by computing phylogenetic and trait resemblance matrices, **S**_**p**_ and **S**_**f**_, based on, phylogenetic tree and species traits, respectively, as shown in Figure 1. The matrices **S**_**p**_ and **S**_**f**_ are standardized by their column totals and multiplied by the transposition of matrix **W** (steps 2.1 and 3.1 in Figure 1) to obtain matrix **P** that represents phylogenetic composition and matrix **X** that represents the functional composition of communities. Computing the square root of Bray-Curtis distance for **P** and **X** we obtain, respectively, **D**_**P**_ and **D**_**f**_ (steps 2.2 and 3.2 in Figure 1), which describes pairwise distances (*D*_*hi*_) among communities (*n*) regarding clade and functional composition. These two distance matrices can be used to obtain the total sum of squares (SS_total_) through Equation 1 and BD_Phy_ and BD_Fun_ by applying Equation 2.

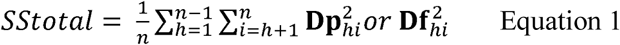

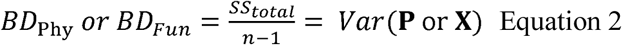

**Figure 1:**
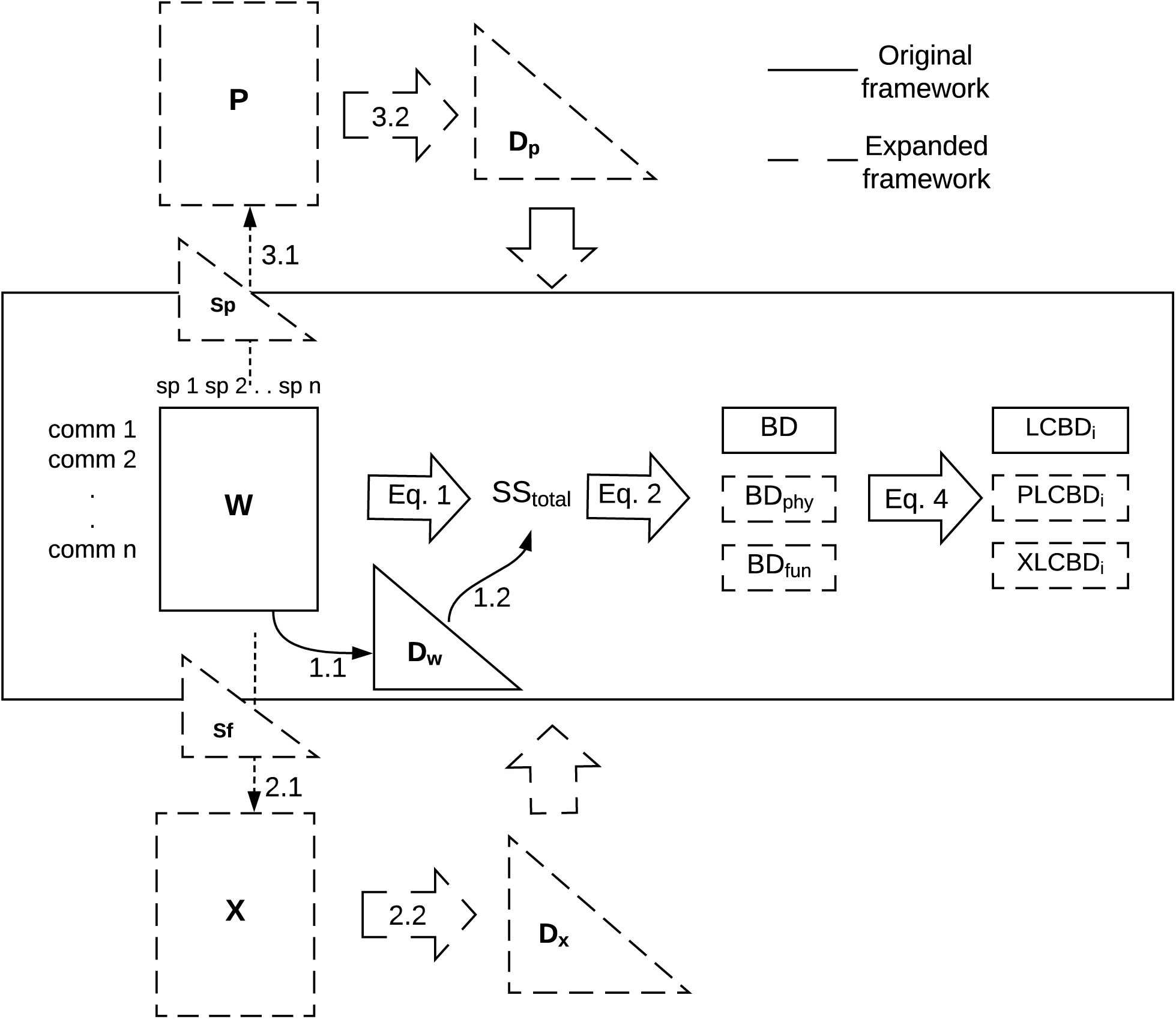
Steps and numerical structures needed extend the BD framework for β-diversity. Solid boxes represent the original numerical structures presented in Legendre & De Cáceres (2013) to calculate BD, while dashed boxes comprise the matrices used in this paper to extend the original framework. The resemblance matrices **S**_**f**_ and **S**_**p**_ are necessary to obtain two matrices that describe communities by their trait (**X**) and phylogenetic (**P**) composition by means of fuzzy weighting. Distance matrices obtained from **P** and **X** are then used to obtain phylogenetic (PLCBD) or trait-based (XLCBD) measures of local contribution for beta diversity, respectively with the equations indicated in arrows.

The components that represents the phylogenetic and functional local contributions of each community, PLCBD and XLCBD respectively, can be obtained by using the algebra of principal coordinate analysis by computing matrix 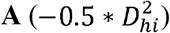 and centering it to obtain matrix **G** through Equation 3. In equation 3, 1 is a vector of ones and 1’ its transposition.

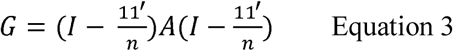

The diagonal elements of matrix **G** are SS_i_ values. Thus, dividing each value in the diagonal of **G** by SS_total_ we can obtain a measure that indicates the proportion that each sample unit accounts for of the total variation presented in **P** or **X** (PLCBD and XLCBD, respectively) (Equation 4).

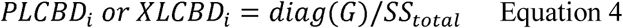

More details on how to decompose BD into its components are provided by Legendre and De Cáceres (2013). The contribution of each species to total β-diversity can only be obtained when using calculations directly on matrices **P** and **X**. Thus, we show here only the components of BD_Phy_ and BD_Fun_ associated with the local contribution of communities. Both PLCBD and XLCBD, like LCBD, can be interpreted as measures of community uniqueness regarding clade and functional composition, respectively. Appendix S1 of the Supplementary Material shows how the BD framework can be extended by directly using matrices **P** and **X** to obtain BD_Phy_ and BD_Fun_, respectively, and all of their components (raw data approach). Figure S1 of the Supplementary Material illustrates how XLCBD and PLCBD can be interpreted using a simple example of a hypothetical metacommunity.

### A simulation-based evaluation of BD_Phy_ and BD_Fun_ as measures of phylogenetic and functional β-diversity

We performed a set of simulations to assess the performance of BD_Phy_, BD_Fun_ and their respective components, PLCBD and XLCBD, in capturing variation in phylogenetic and functional structure of metacommunities. The simulation procedure is based on the protocol proposed by Peres-Neto et al. (2012) (see also Minchin, 1987 for the original simulation approach), and allows the integration of phylogenetic relationships, species traits, species composition and environmental gradients in different ways to determine if the distribution of attributes of species and clades across communities mediate variation in metacommunity composition in a simulated environmental gradient. The procedure starts with a simulation of a phylogenetic tree and species traits. The presence of a species in a community is determined by a probability function that corresponds to a match between a species trait value and a simulated environmental value for each community.

We simulated four scenarios to test the performance (type I error and power) of the new metrics obtained using the expanded framework for β-diversity: (1) metacommunities in which phylogenetic and functional composition are responsible for variation in metacommunity composition (scenario W1,P1,X1); (2) metacommunities in which only clade composition is responsible for variation in metacommunity composition (scenario W1,P1,X0); (3) metacommunities in which only functional composition is responsible for variation in metacommunity composition (scenario W1,P0,X1); and, finally, (4) metacommunities in which neither clade distribution nor functional composition are responsible for variation in metacommunity composition (scenario W1,P0,X0).

To generate metacommunities according to scenario W1,P1,X1, we simulated a phylogenetic tree in which species presented traits with high phylogenetic signal, so that both trait and phylogenetic composition will vary across the metacommunity. In scenario W1,P1,X0, metacommunities were assembled from a phylogenetic tree in which species traits have phylogenetic signal, however, the trait used to calculate BD_Fun_ and XLCBD was not the same as the one used to assemble the metacommunity, thus, phylogenetic composition must show variation across the metacommunity, but not functional composition. In scenario W1,P0,X1, the metacommunities were assembled from a phylogenetic tree in which species did not present traits with phylogenetic signal, thus, functional composition will vary across the metacommunity, but not phylogenetic composition. Finally, in scenario W1,P0,X0, the metacommunities were assembled from a phylogenetic tree in which species traits did not present phylogenetic signal and the trait used to calculate BD_Fun_ and XLCBD was not the same as the one used to assemble the metacommunities, thus, neither functional nor distribution of clades were responsible for variation in composition across the metacommunity.

All metacommunities were simulated to possess 50 communities with species selected from a pool of 200. The simulation procedure was repeated 999 times for each scenario. Type I error and power for BD_Fun_, BD_Phy_, PLCBD and XLCBD were calculated by counting the number of times that the null hypothesis was rejected in each round of the simulation procedure. Each round consisted of: (1) simulating the phylogenetic tree, species traits and metacommunities; (2) calculating BD_Phy_, BD_Fun_ and the local components PLCBD and XLCBD; (3) running a null model that deconstructs the original phylogenetic and functional relationships among species (taxa shuffle null model, Kembel et al., 2013) and recalculating BD_Phy_, BD_Fun_, PLCBD and XLCBD; (4) repeating step 3 three 999 times to assemble a null distribution of metrics; and (5) comparing the observed values of BD_Phy_ and BD_Fun_ with the null distributions of BD_Phy_ and BD_Fun_ using α = 0.05 as the nominal error to reject the null hypothesis. To test the performance of the metrics of local contribution, in step 4 we also obtained a F statistic from a linear model that related observed PLCBD and XLCBD with the environmental gradient used in the simulation and compared observed F with a null F distribution obtained from a linear model that relates the null PLCBD and null XLCBD with the environmental gradient. We compared the null F distribution with the observed F, also using the nominal α = 0.05 to reject the null hypothesis (illustration of performance analysis is shown in Figure S3 of Appendix 2). More details on simulation procedures, models and theoretical expectations regarding PLCBD and XLCBD are presented in Appendix S2.

A summary of the four scenarios used to test the performance of the metrics proposed in this work is provided in Figure 2. Dimensions that presented variation in metacommunity composition in the simulation process are represented by the number one, while dimensions that did not mediate variation in the simulation process of the metacommunity are represented by a zero. Figure 2 also summarizes which scenarios were used to test power and which were used to test type I error. We did not perform tests of the original BD and LCBD since the statistical behaviors of these metrics were previously tested by Legendre and De Cáceres (2013). For more details about the simulation procedure and methods to obtain F values (models used to assess the performance of PLCBD and XLCBD) see Appendix S2 in Supplementary Material. We also tested the performance of all metrics proposed in this work using the raw data approach (Supplementary material).

**Figure 2:**
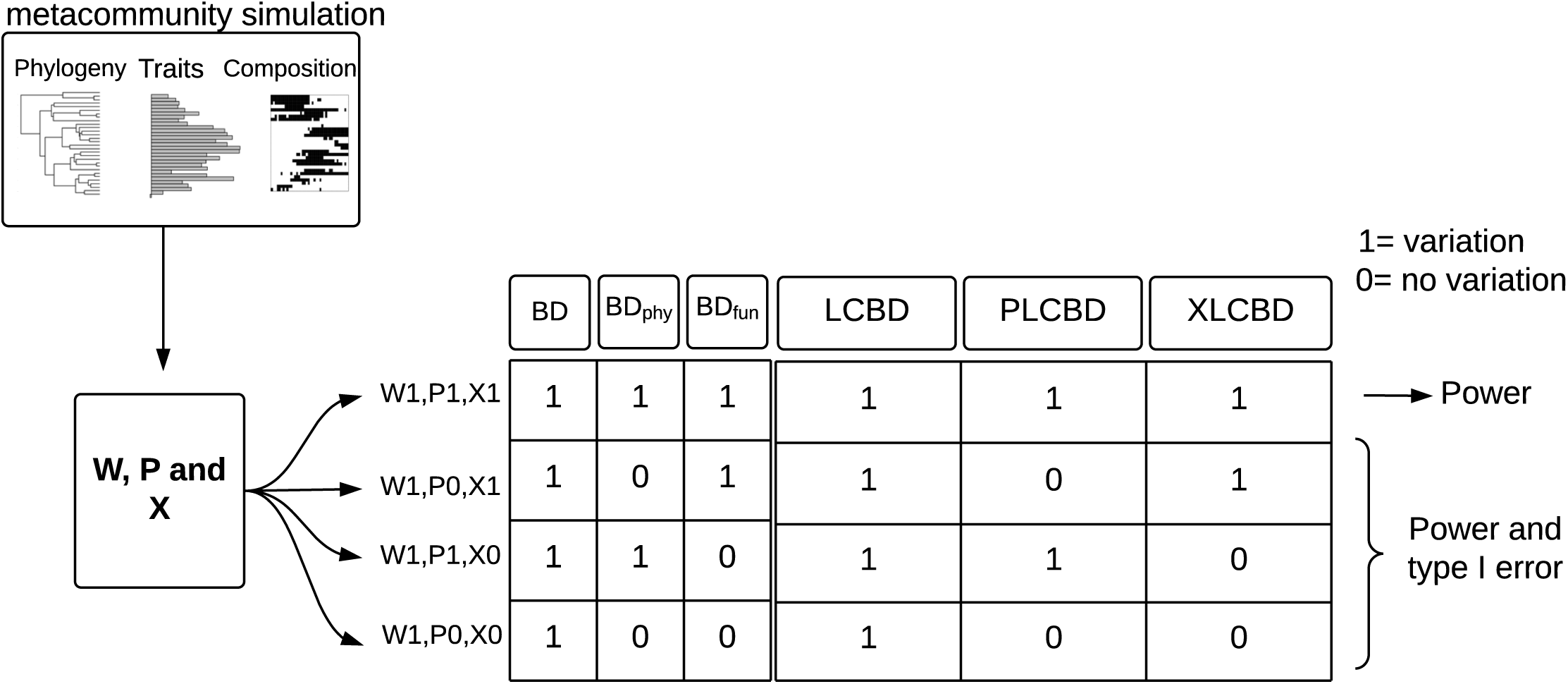
Schematic representation of all simulation scenarios used to test the metrics proposed in this work. **W** represents a metacommunity that is generated by a simulation process, **P** and **X** represents functional and phylogenetic structure of metacommunity. When a given component of diversity presented variation in the metacommunity we assigned the number 1, on the contrary we attribute 0 in the table. The performance tested with each one of the scenarios (type I, power or both) is specified beside each scenario.

### Empirical application: β-diversity of stream fish communities of a tropical river basin

We illustrate how the new metrics proposed here can be used by calculating BD_Phy_, BD_Fun_, PLCBD and XLCBD for tropical stream fish communities located in Brazil. This data comprises a sample of a metacommunity with 173 sites located in the Ivinhema River Basin, which is the main tributary of the Paraná River in the western part of the Paraná River Basin, one of the largest basins in South America. Each site corresponded to a stretch of approximately 100 meters in length and was sampled with the aid of an 80 x 120-cm rectangular sieve with a mesh size of 2mm. Electrofishing was also used at some sites.

We calculated BD, BD_Phy_, BD_Fun_ and their local components, LCBD, PLCBD and XLCBD, for the 173 sampled sites. The BD metric was calculated using an incidence matrix standardized by the total occurrence of species in a community and subjected to a chord transformation. This standardization was done in order to keep BD, BD_Phy_ and BD_Fun_ on the same scale of variation. We assessed the significance of these metrics using two null models: site shuffle and taxa shuffle. Site shuffle randomizes species occurrence in the metacommunity by shuffling the lines of matrix **W**, with the rejection of the site shuffle null model (α ≤ 0.05) indicating that the composition of the community differs from that expected by chance. The taxa shuffle null model randomizes the tips of the phylogeny and functional dendrogram and uses these randomized structures to calculate, respectively, matrices **P** and **X**. The rejection of taxa shuffle indicates that the uniqueness of a community is also due to deep evolutionary relationships among the species or their functional attributes.

We performed a correlation analysis among LCBD, PLCBD and XLCBD using Pearson’s correlation index, which allowed us to assess which dimensions serve as a proxy to indicate the local contribution of other dimensions. Finally, we represent the three quantities of local contribution spatially through an RGB plot to show how these three components can be used together to identify sites of high importance regarding uniqueness. To generate the RGB plot we first standardize LCBD, PLCBD and XLCBD to vary between 0-250 and attribute to each combination of the three metrics a color in the gradient of RGB system. The redder the color the greater the uniqueness of a community regarding taxonomic composition; the greener the color the greater uniqueness regarding clade composition; the bluer the color the greater the uniqueness regarding functional composition. We also calculated how much each species contributed to β-diversity of Ivinhema River Basin regarding taxonomic (SCBD), phylogenetic (PSCBD) and functional (XSCBD) components (Appendix S4).

More details regarding the sampling design, the phylogenetic hypothesis and traits of the fish metacommunity used to calculate the metrics are provided in Nakamura et al. (2017) and Appendix S4. In Appendix S5 in Supplementary Material we provide the R function used to calculate all the metrics proposed in this work and for testing their significance according to the taxa shuffle null model (Kembel et al. 2013). We suggest that the distance metric to be used for the two metrics proposed here be the square root of Bray-Curtis index, which produces a maximum value of 0.5 for BD_Phy_ and BD_Fun_ for communities with completely different compositions. For the raw data procedure, we suggest the use of chord transformation of matrices **P** and **X** prior to the calculation of BD_Phy_ and BD_Fun_ metrics, which will produce values ranging from 0 to 1.

## Results

### Performance of metrics

We present here only the results of the simulation analysis for metrics calculated with the distance-based approach, while the results for the raw data approach are provided in Table S1 of Appendix S3 in Supplementary Material. **Error! Reference source not found.** shows the statistical performance (rejection rate, type I error and power) for BD_Phy_, BD_Fun_, PLCBD and XLCBD, and the mean R^2^ of the linear models relating the simulated environmental gradient to the metrics PLCBD and XLCBD, calculated for the four simulated scenarios. Both BD_Phy_ and BD_Fun_ had acceptable type I error values for all scenarios, with BD_Phy_ having a value of 0.05 for both W1,P0,X0 and W1,P0,X1, and BD_Fun_ presenting a probability of 0.04 for both W1,P0,X0, and W1,P1,X0. Similar results were found for the raw data approach, in which BD_Phy_ had values of 0.04 and 0.05 in scenarios W1,X0,P0 and W1,X1,P1, respectively, whereas BD_Fun_ had values of 0.05 for both W1,X0,P0 and W1,X0,P1(Table S1 of Appendix S3 in Supplementary Material). Both BD_Phy_ and BD_Fun_ presented high values for power for all scenarios tested (ranging from 0.99 to 1).

PLCBD had a type I error rate of 0.04 in both W1,P0,X0 and W1,P0,X1, whereas XLCBD had type I error rates of 0.04 and 0.05 for W1,P0,X0 and W1,P1,X0, respectively. Both PLCBD and XLCBD had high rates of power ranging from 0.66 to 1. The lowest power value was obtained for PLCBD in scenario W1,P1,X1 for both the distance-based and raw data approaches (0.66 for both).

### Application of extended framework to stream fish communities

BD, BD_Phy_ and BD_Fun_ presented values of, respectively, 0.38, 0.09 and 0.01 for the stream fish metacommunity of Ivinhema River Basin. BD_Fun_ and BD_Phy_ did not differ significantly from expected values of the taxa shuffle null model (p-values > 0.05). The contribution of local communities ranged from 0.003 to 0.009 for LCBD and 0.002 to 0.02 for both PLCBD and XLCBD. Regarding uniqueness in composition, 23 communities presented significant values for the site shuffle null model for LCBD, 26 for PLCBD and 45 for XLCBD. Of the 26 significant values for PLCBD, 11 were also significant for taxa shuffle, indicating the effects of deep evolutionary relationships in phylogenetic uniqueness of these communities. For XLCBD, 15 communities (from the 45 that presented significant p-values for site shuffle) presented significant values for the taxa shuffle null model. Communities that presented significant values for PLCBD and XLCBD for site and taxa shuffle are represented in Figure 3 by, respectively, crossed circles and triangles.

**Figure 3:**
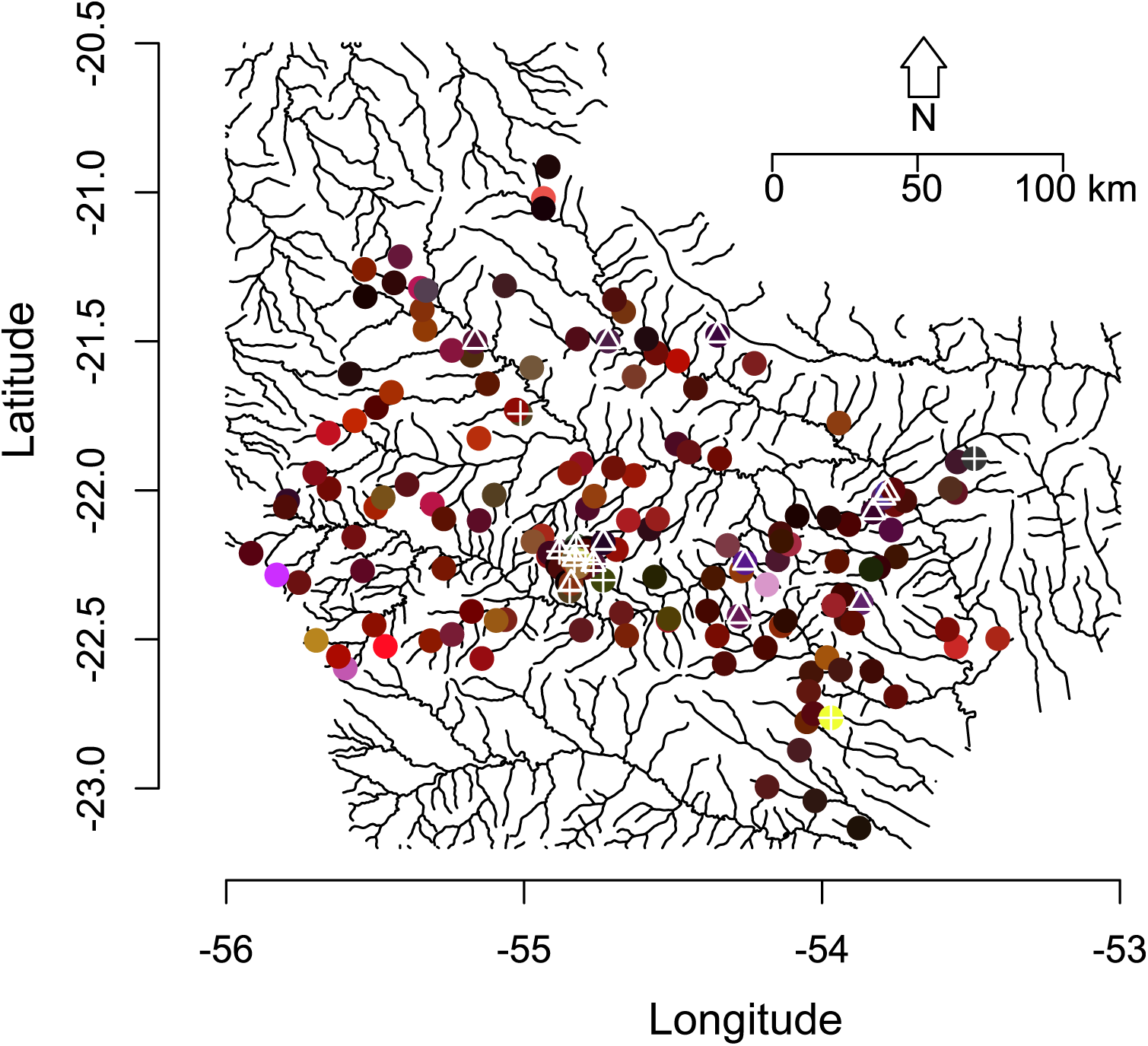
Map showing the distribution of LCBD, PLCBD and XLCBD values for fish communities of Ivinhema River Basin. Each point represents a combination of values of local contribution for the three dimensions. The redder the color the greater the uniqueness of a community regarding taxonomic composition; the greener the color the greater uniqueness regarding clade composition; the bluer the color the greater the uniqueness regarding functional composition. Communities that presented significant values for PLCBD and XLCBD for site and taxa shuffle are represented by points with, respectively, crossed circles and triangles.

All local components presented low correlation among each other, with the highest correlation found being for LCBD and PLCBD (ρ= 0.46; p-value < 0.001). This lack of correspondence among components of local contribution can also be noted in Figure 3. Figure 3 shows a predominance of a combination of low values for PLCBD, XLCBD and PLCBD (circles with black color), indicating the predominance of communities with low uniqueness. Only a few communities presented high values for at least one component, with LCBD being the component with more communities with high uniqueness regarding taxonomic composition (represented by red circles). The presence of high values for two components is even more rare among the communities analyzed, indicating that is very rare for the communities of Ivinhema River Basin to host unique species, clades or species that present very distinct attributes. In Appendix S4 we also show the contribution of each species for Beta diversity of metacommunity.

## Discussion

We demonstrated that the BD framework proposed by Legendre and De Cáceres (2013) can be effectively extended to accommodate other components of biodiversity. The extension presented here incorporates the advantages of the original BD framework (decomposition into meaningful components of local and species contributions) while at the same time effectively assessing β-diversity for phylogenetic and functional dimensions of biodiversity, as shown by the performance analysis of the metrics.

As far as we are aware, the extension proposed in this work is the most general in literature, since other propositions that seek to expand Legendre and De Cácereś framework lack some important characteristics presented in the original proposition. Shooner et al. (2018) proposed a phylogenetic informed LCBD (also named as PLCBD) by using a phylogenetic distance matrix containing PhyloSor values (Bryant et al. 2009) for calculating LCBD. Since the use of a phylogenetic distance matrix among sites only allows BD to be decomposed into the local contribution component, it is not possible to obtain the portion of BD that accounts for species contribution to the total variation of the metacommunity (SCBD component). The use of the fuzzy weighting method together with Legendre and De Cácereś equations allows phylogenetic and functional informed BD and all of its components to be derived, since the calculations can be made using both distance and raw data approaches (using directly matrix **P** or **X**). Besides the generality presented in our extension, we also highlight its flexibility, to obtaining new β-diversity measures that capture other dimensions of biological diversity. For example, a matrix containing genetic distances among species can be used in fuzzy-weight transformation to obtain the genetic composition of sites (e.g. see Duarte et al., 2018), which in turn can be used to compute a genetic informed BD measure and its components. Therefore, the method presented here can be viewed as a unified approach that allows the derivation of a family of β-diversity metrics with the same mathematical characteristics yet encompassing different components of biological diversity.

The new local component metrics, PLCBD and XLCBD, can provide interesting tools to address questions in the investigation of patterns of organization of metacommunities. PLCBD can be used to identify patterns of variation present in the phylogenetic structure of metacommunities related to sites that host clades with unique evolutionary history. This information can be useful to test hypothesis that seek to understand how the degree of phylogenetic uniqueness of sites is related to environmental or historical factors (Graham & Fine, 2008; Leibold et al, 2010). Other methods, like PCPS (Duarte, 2011), can also be used to identify distinct sites regarding phylogenetic composition (heuristically, accompanied by an ordination procedure), however, PLCBD offers a more direct assessment of community uniqueness together with hypothesis testing to evaluate the role of species composition and deep evolutionary relationships in generating observed patterns of phylogenetic uniqueness (through site and taxa shuffle null models).

Regarding practical applications of PLCBD and XLCBD, we highlight their utility for conservation purposes by identifying sites that deserve special attention due to uniqueness in species composition, evolutionary history and functional attributes. In this way they serve as complementary measures for studies that use local uniqueness based solely from a taxonomic perspective (e.g. Landeiro et al. 2018). An integrative approach that addresses multiple dimensions of variation in biological diversity can be used to identify areas of congruence or mismatches in the distribution of biological diversity, which can influence decisions made in conservation plans to preserve regional biotas (Devictor et al. 2010; Meynard et al. 2011).

Our empirical example using a stream fish metacommunity illustrates how to link patterns of variation in β-diversity related to taxonomic, phylogenetic and functional diversity. Our findings complement those of Súarez et al (2011), by evidencing that the processes influencing the distribution of species in the upper Ivinhema River Basin are mainly mediated by variation in species composition and contemporary environmental factors, with the deep evolutionary history of the species having less impact on the structure of this metacommunity. By mapping LCBD, PLCBD and XLCBD we can see that only a few sites possess unique compositions of species, clades and attributes. Our multifaceted approach for beta diversity illustrates how these three metrics can be used together to help identify areas of special conservation interest (e.g., Devictor et al. 2010; Landeiro et al. 2018). The lack of a spatial pattern in local contribution indicates that the communities have very similar importance for the maintenance of the entire β-diversity of the Ivinhema River Basin.

### Concluding remarks and future directions

We move forward in the operationalization of β-diversity patterns by presenting a simple way to extend the BD framework in order to derive effective phylogenetic and functional β-diversity measures. The methods used in this work to extend the original framework are general enough to be used as a basis to obtain other metrics that can represent other dimensions of biological diversity while at the same time preserve the unique advantages presented in the original BD framework.

## Authors’ declaration

GN and LD conceived the idea, performed the analysis and wrote manuscript. WV and YRS collected the data for stream fish communities and species traits, and constructed the phylogenetic hypothesis used in this study.

## Acknowledgements

Research of LD and GN was conducted in the context of the National Institute for Science and Technology (INCT) in Ecology, Evolution and Biodiversity Conservation, supported by MCTIC/CNPq (proc. 465610/2014-5) and FAPEG (proc. 201810267000023). Research activities of LD was supported by a CNPq Productivity Fellowship (grant 307527/2018-2).

