## Supplementary material for "A multifaceted approach to analyzing taxonomic, functional and phylogenetic β-diversity"

Appendix S1

*Obtaining BD through the raw data approach*

We summarize here how BD_Phy_ and BD_Fun_ can be obtained using the raw data approach. The major advantage of calculating BD metrics this way is that it allows BD to be decomposed into a component that represents species contribution to beta diversity (SCBD). We start with a matrix that describes phylogenetic or functional composition of communities (matrix **P** or **X,** respectively) obtained using a fuzzy weighting procedure and use it to compute the square deviation of the proportion of incidence of each species in a community (y_ij_) with its respective mean incidence in the community ( ̅yj) as described in Equation S1. Each value obtained from Equation S1 composes matrix **S** of square deviations.

$s_{ij}= \left( y_{ij}- \bar{y_{j}} \right)^{2}$ Equation S1.

Using Equation S2 we can obtain the BD measure, where *n* corresponds to the total number of sites in the metacommunity and *p* the total number of species.

${BD}_{Phy} or {BD}_{Fun}=\frac{\sum_{i=1}^{n} \sum_{j=1}^{p} s_{ij}}{n-1}$ Equation S2

Since *BD_Phy_* and *BD_Fun_* comprise the total variation in, respectively, matrix **P** and **X**, we can partition it into different components that represent the variation accounted for by the rows of matrix **S** and the variation accounted for by the columns. By summing the rows of matrix **S** we obtain a measure of local contribution for beta diversity (PLCBD for phylogenetic contribution and XLCBD for functional contribution) and by summing the elements in each column of matrix **S** we obtain a measure of species contribution for beta diversity (PSCBD for phylogenetic and XSCBD for functional).

*Interpreting phylogenetic and functional local components of BD_Fun_ and BD_Phy_*

In order to clarify how components PLCBD and XLCBD can be interpreted as measures of phylogenetic and functional uniqueness, we illustrate in Figure S1 a hypothetical example with a metacommunity. The metacommunity is described by the presence of five species (matrix **W**) with their respective phylogenetic relationships (phylogenetic tree) and a functional dendrogram describing trait similarities among species. Three species occur only in community 1 (sp 2, sp 3 and sp 5 in matrix **W**), consequently, community 1 has the highest value for LCBD, since this is the most unique in terms of species composition. On the other hand, community 3 hosts only species 1, and is the most distinct in terms of evolutionary history. For this reason, despite the lower contribution to the taxonomic component of beta diversity (low LCBD), community 3 has the highest PLCBD (0.50). Finally, community 2 is the most unique in terms of functional composition since it hosts species 4, and is the most distinct regarding its attributes (illustrated in the functional dendrogram).

Figure S1: Schematic representation of a simple example containing one metacommunity composed of three communities and five species, with their occurrence and phylogenetic and functional relatedness described by, respectively, matrix **W**, a phylogenetic tree and a functional dendrogram. Matrices **X** and **P** are the results of a fuzzy weighting procedure applied to matrix **W**, and now describe functional and phylogenetic composition, respectively. Distance matrices **D_x_**, **D_w_** and **D_p_** are calculated according to an appropriate dissimilarity index and describe dissimilarity among communities according to functional, taxonomic and phylogenetic dimensions. A measure of the contribution of each local component to total β-diversity related to functional (XLCBD), taxonomic (LCBD) and phylogenetic (PLCBD) dimensions of diversity can be obtained using Equation 4.

Appendix 2

*Simulation procedure.*

To simulate the four scenarios used to test the performance of BD_Phy_, BD_Fun_ and their respective component metrics PLCBD and XLCBD, we use an adaptation of the simulation procedure proposed by Peres-Neto et al.(2012), which in turn is an adaptation of a procedure detailed by Minchin (1986) to simulate community composition in n-dimensional environmental gradients. The simulation procedure can be viewed as a three-step process that comprises the following steps: (1) simulation of a phylogenetic tree that describes the evolutionary relationships among species that will compose the species pool; (2) simulation of the traits of these species; and (3) the assembly of a metacommunity based on the traits and a simulated environmental gradient used to allocate species among local communities. Each community in the metacommunity represents a point in the environmental gradient. Below we detail each of these three steps used to simulate metacommunities.

*1 – Simulation of the phylogenetic tree.*

To simulate the phylogenetic tree we used the function *sim.bdtree* of the *geiger* package (Harmon et al., 2008), which simulates the relatedness of species by a homogeneous birth-death process across all lineages. We set the speciation rate to 0.1 and the extinction rate to zero. This phylogeny was then used to simulate traits for the species.

*2 – Simulation of species traits.*

We simulated traits using the function *rTraitCont*. We manipulated the phylogenetic signal of traits by modifying the branch lengths of the phylogenetic tree using Grafen’s ρ parameter and the function *compute.brlen.* Both *rTraitCont* and *compute.brlen* are of the *ape* package (Paradis et al., 2004). Low values for parameter ρ lengthen shallow branches of the tree, simulating ancient diversification of species, and thus generate low phylogenetic signal in traits. On the other hand, values of ρ that are near 1 lengthen deeper branches, simulating trait variation equivalent to that expected by a Brownian motion model, thus generating high phylogenetic signal in traits. We used ρ = 0.0001 to simulate traits with low phylogenetic signal and ρ = 1 to generate traits with high phylogenetic signal. Trait values were scaled to vary between -1 and 101 prior to subsequent analyses.

This trait simulation procedure was previously tested by Duarte et al. (2016), who demonstrated that modifications of ρ generated phylogenetic signal as expected; i.e., low values of ρ generating low phylogenetic signal and values of ρ = 1 generating phylogenetic signal near 1, as quantified by the K statistic (Blomberg, Garland Jr, & Ives, 2003).

*3- Metacommunity assembly*

From the simulated phylogenetic tree and traits (referred as **F**), we simulated metacommunities using a function that relates the values of attributes in vector **F** with values of an environmental gradient **E** present in each community (with each value in these vectors represented as *f_i_* and *e_k_*, respectivelly). The values of the environmental gradient were simulated from a uniform distribution ranging from 0 to 100. Species abundances in communities were simulated as unimodal response curves where the abundance w*_ik_* of the *i*^th^ species in the *k*^th^ community (Peres-Neto et al. 2012) is defined as follows:

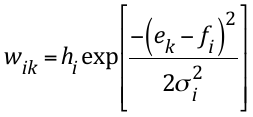
Equation 1

Vector **h** in equation 1 contains random uniformly-distributed values ranging from 0 to 30 and represents the maximum abundance of each species at its optimum. Based on Equation 1 we simulated metacommunities containing 50 communities.

In summary, the simulation procedure described in Equation 1 can be interpreted as a sampling procedure of each species across an environmental gradient such that each species has distinct probabilities of being sampled at different locations along this gradient (Figure S1). These differences guarantee that the simulated metacommunity would possess variation in the species composition of its communities.

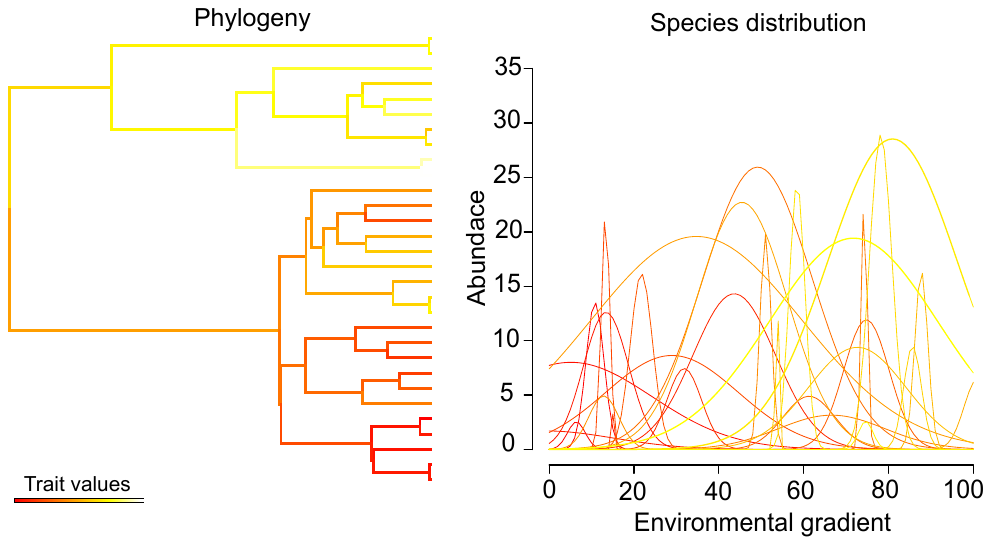

Figure S2: Schematic representation of a phylogenetic tree with species traits represented by colors and a graphic of unimodal response curves for each species present in the phylogeny. Similar species regarding their traits and phylogenetic relationships have a greater probability of being sampled in communities possessing similar values along the environmental gradient. This example illustrates the unimodal response curves for one metacommunity created according to scenario W1,P1,X1.

The four scenarios used to test the metrics proposed in this work were produced by combining the presence or absence of phylogenetic signal in traits with the traits used to calculate the metrics in the following manner: For the situation in which phylogenetic relationships and attributes of species mediate variation in community composition across the metacommunity, species that present similar traits and are phylogenetically similar have greater probability of being sampled at similar points along the environmental gradient. This situation corresponds to the scenario W1,P1,X1, which is illustrated in Figure S2 as an example of the format of unimodal species response curves. For the situation in which variation in community composition is mediated only by the attributes of species, the traits used in assembling the metacommunity do not possess phylogenetic signal, which corresponds to scenario W1,P0,X1. If only phylogenetic relationships mediate variation in community composition, then the traits used to calculate BD_Fun_ are not the same as those that mediate metacommunity assembly (scenario W1,P1,X0). If neither attributes nor phylogenetic relationships of species mediate variation in community composition, the traits used to calculate BD_Fun_ are not the same as those that are responsible for metacommunity assembly, and those that were used to simulate the metacommunity do not possess phylogenetic signal (W1,P0,X0).

The theoretical expectation regarding PLCBD and XLCBD are illustrated in Figure 3. Metacommunities structured by traits with phylogenetic signal will present higher values of PLCBD and XLCBD for communities near the extremes of environmental conditions than communities with environmental values near the mean for the entire metacommunity (high observed F values than that generated by the null models). On the other hand, metacommunities in which the assembly process is not dependent on the phylogenetic relationships of species nor by the traits they possess, will present similar values of PLCBD and XLCBD across all the entire environmental gradient (observed F values similar to that generated by null models).

Figure S3: Schematic representation of simulation procedure used to test the performance of PLCBD and XLCBD. Scenario A represents a metacommunity simulated to test the power of PLCBD and XLCBD, in which communities in the extreme of the environmental gradient will present higher values of PLCBD and XLCBD than communities in the middle of the gradient. As a consequence, the F values derived from a model relating PLCBD/XLCBD with environment (F_obs_) to simulate the communities must be higher than the F values generated from a null model that related null PLCBD/XLCBD. In B we present a metacommunity simulated to test the type I error rate of PLCBD/XLCBD. In this situation F_obs_ must not be different from a null distribution of F values derived from a null model (using as critical value α=0.05).

Appendix S3

Table S1 Provides the results for the raw data approach for calculating BD_Phy_, BD_Fun_ and their respective components PLCBD and XLCBD.

Table S1: Statistical performance, type I error (alpha 0.05) and power of BD_Phy_, BD_Fun_ and their components PLCBD and XLCBD calculated by the raw data approach.

| Scenario | BD_Phy_ | BD_Fun_ | PLCBD | | | | | XLCBD | | | | |
| --- | --- | --- | --- | --- | --- | --- | --- | --- | --- | --- | --- | --- |
|  | **Rejection rate** | **Rejection rate** | **Mean R2** | **TI site** | **TI taxa** | **Pw site** | **Pw taxa** | **Mean R2** | **TI site** | **TI taxa** | **Pw site** | **Pw taxa** |
| W1,P0,X0 | 0.04 | 0.05 | 0.44±0.18 | 0.94 | 0.04 | - | - | 0.25±0.15 | 0.82 | 0.03 | - | - |
| W1,P1,X0 | 0.98 | 0.05 | 0.78±0.16 | - | - | 0.98 | 0.80 | 0.22±0.17 | 0.67 | 0.05 | - | - |
| W1,P0,X1 | 0.05 | 1 | 0.45±0.17 | 0.95 | 0.03 | - | - | 0.90±0.04 | - | - | 1 | 1 |
| W1,P1,X1 | 0.96 | 1 | 0.53±0.14 | - | - | 0.98 | 0.66 | 0.60±0.10 | - | - | 1 | 0.96 |

Scenario = different scenarios used to simulate metacommunities; TI site = type I error for site shuffle procedure; TI taxa = type I error for taxa shuffle procedure; Pw site = statistical power for site shuffle procedure; Pw taxa = statistical power for taxa shuffle procedure.

Appendix S4

*Phylogenetic hypothesis and species traits*

We constructed a phylogenetic tree describing the evolutionary relationship among fish species of Ivinhema River Basin using as a backbone phylogeny the hypothesis proposed by Betancur-R *et al*. (2013). We used other works with specific groups (Santos, 2007; Oliveira *et al*. 2011; Nakatani *et al*. 2011; Armbruster, 2004; Sullivan et al. 2013; Lavoué *et al*. 2012; Hertwig, 2007) to complement and improve the resolution of some families present in the backbone phylogeny. Species present in the sampled sites, but absent in the phylogeny. were replaced by the phylogenetically most closely related species available in the Betacur-R et al. (2013) phylogeny. Species with poorly established phylogenetic relationships, and for which we could not find any study to help elucidate their position, were included at family level, which generated some polytomies in the tree. The phylogeny was data by applying the BLADJ (Branch Length Adjustment) algorithm in Phylocom software (Webb et al. 2008), which estimates the length of non-dated nodes evenly through dated nodes. Dated nodes were those provided by Betancur-R et al. (2013). The phylogeny used to obtain cophenetic distance among species, dated in million years, is presented in Figure S4

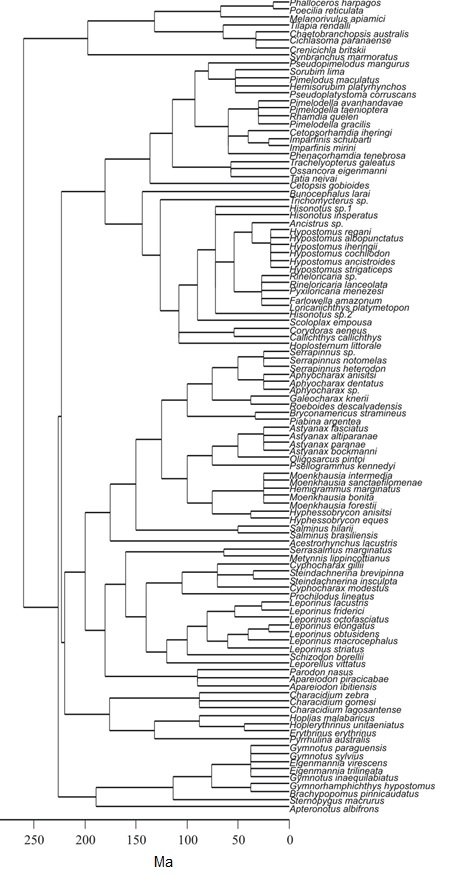

Figure S4: Phylogenetic hypothesis used to obtain cophenetic distance among stream fish species from Ivinhema River Basin, dated in million years.

We adopted a broad definition of functional traits as being those attributes of species that directly or indirectly influenc species performance (*e.g*. growth, reproduction, mortality). In summary, we selected a set of functional traits that are related to feeding, habitat occupation and life history (Gatz Jr. 1979; Winemiller, 1991; Oliveira, 2005). Some linear measures and areas of morphological structures were taken on the left side of three to five adult individuals in order to avoid trait differences do to ontogenetic variation in traits. Others measures (trophic level and maximum length) we obtained from Fishbase (Froese and Pauly, 2013), an online database for fishes. All measures used in this work along with their characteristics and definitions are summarized in Table S2.

Table S2: Functional attributes used to obtain functional distance for fish communities of Ivinhema River Basin

| Functional attribute/ Ecomorphological index | Definition | Functional category | Type (continuous or categorical) |
| --- | --- | --- | --- |
| Maximum length (ML) | Maximum length known for the specie or the length of the longest specimen sampled. Measured as the distance between the anterior part of the head and the end of the caudal fin. | Habitat use, life history characteristics and diet. | continuous |
| Relative eye area (REA) | Ratio between the horizontal distance between margin of eyes and squared maximum length. High values are associated with species with greater visual capacity and that inhabit upper areas in the water column (Gatz Jr., 1979). | Diet, habitat use. | continuous |
| Vertical position of eye (VPE) | Ratio between the vertical distance to the center of pupil and height of head. This measure is associated with foraging position of species in the water column. High values indicate benthonic fishes, low values indicate nektonic fishes. (Gatz Jr., 1979). | Habitat use | continuous |
| Relative height (RH) | Ratio between maximum vertical distance and maximum length. Inversely related with water velocity and related to ascendant and descendant movements (Gatz Jr., 1979; Winemiller, 1991). | Locomotion, habitat use. | continuous |
| Aspect-ratio caudal fin (ACF) | Ratio between the squared maximum vertical distance of caudal fin and area of caudal fin. High values indicate greater swimming capacity (Gatz Jr., 1979; Breda, Oliveira & Goulart, 2005). | Locomotion. | continuous |
| Relative caudal peduncle length (CPL) | Ratio between distance from posterior proximal margin of anal fin and maximum length. Longer peduncles indicate species adapted to rifles (Oliveira, 2005). | Locomotion. | continuous |
| Relative mouth width (MW) | Ratio between horizontal distance measured inside of fully open mouth at widest point and maximum length. Greater values indicate species with preferences for larger prey (Gatz Jr., 1979). | Diet. | continuous |
| Trophic level (TL) | Species position in the food web, estimated as 1 plus the mean of trophic level of food items in the diet, weighted by the contribution of different food items. | Diet. | categorical |

*Taxonomic, functional and phylogenetic contribution of species*

Calculation with the raw data approach allows BD to be decomposed into one additional source of variation — species contribution to beta diversity (SCBD). We calculated the original SCBD for species of Ivinhema River Basin, as well as the phylogenetic and functional SCBD (namely PSCBD and XSCBD, respectively). Figure S3 illustrates the rank of species contribution relative to these three components of biodiversity. Each dot represents a species, species names in each graphic represent the three most important species to total BD.

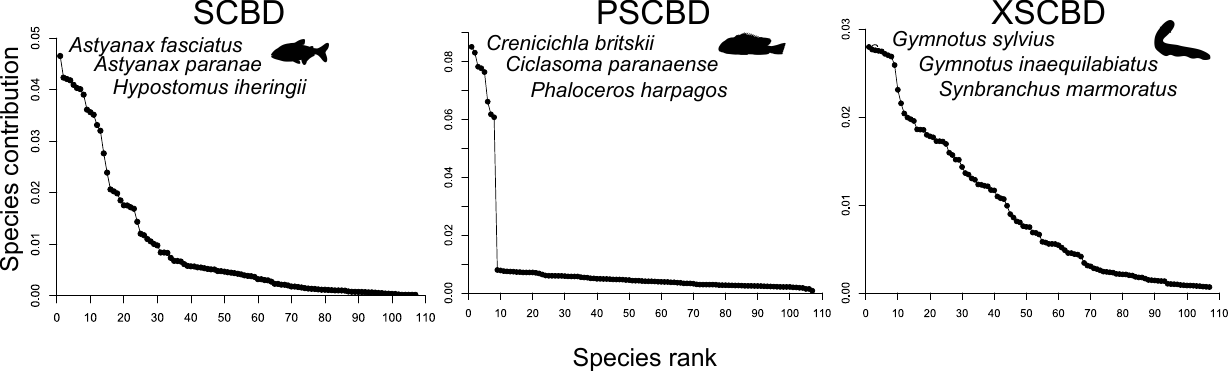

Figure S4: Species rank based on their contribution to BD, BD_Phy_ and BS_Fun_. Each dot represents a species and the names in each graphic represent the three most important species. The silhouette is a representative of each group of most important species.

Appendix S5

We provide an R function that extends the calculation of BD to include phylogenetic and functional dimensions of diversity using raw data and distance-based approaches, this function and all the updates can be found at https://github.com/GabrielNakamura/BetaDiv_extension. The function calculates the original metric proposed by Legendre and Cáceres (2015), called here WLCBD, and the two new metrics proposed in the present study, PLCBD and XLCBD for phylogenetic and functional dimensions, respectively. We also provide in the function two null models to test the significance of the contribution of each community to the three components of diversity. The first null model, called site shuffle, tests the null hypothesis that the local contribution is a result of random community structure. The second null model, called taxa shuffle, tests the null hypothesis that the local contribution of communities for the functional and phylogenetic dimensions is the result of random phylogenetic or functional structure.

Rejection of the first null hypothesis (p<0.05) generated by site-shuffle, indicates that the contribution of a site to beta diversity is a result of a non-random mechanism that structures the observed community composition (species co-occurring in sites). Rejection of the second null hypothesis (p<0.05) generated by taxa shuffle, indicates that the local contribution to functional and phylogenetic diversity is the result of functional or phylogenetic similarities observed between species.
